# Neural decoding of bistable sounds reveals an effect of intention on perceptual organization

**DOI:** 10.1101/206417

**Authors:** Alexander J. Billig, Matthew H. Davis, Robert P. Carlyon

**Author notes:** Corresponding Author: Alexander J. Billig, UCL Ear Institute, 332 Gray’s Inn Road, London, WC1X 8EE, United Kingdom.

## Abstract

Auditory signals arrive at the ear as a mixture that the brain must decompose into distinct sources, based to a large extent on acoustic properties of the sounds. An important question concerns whether listeners have voluntary control over how many sources they perceive. This has been studied using pure tones H and L presented in the repeating pattern HLH-HLH-, which can form a bistable percept, heard either as an integrated whole (HLH-) or as segregated into high (H-H-) and low (-L—) sequences. Although instructing listeners to try to integrate or segregate sounds affects reports of what they hear, this could reflect a response bias rather than a perceptual effect. We had human listeners (15 males, 12 females) continuously report their perception of such sequences and recorded neural activity using magneto-encephalography. During neutral listening, a classifier trained on patterns of neural activity distinguished between periods of integrated and segregated perception. In other conditions, participants tried to influence their perception by allocating attention either to the whole sequence, or to a subset of the sounds. They reported hearing the desired percept for a greater proportion of time than when listening neutrally. Critically, neural activity supported these reports; stimulus-locked brain responses in auditory cortex were more likely to resemble the signature of segregation when participants tried to hear segregation than when attempting to perceive integration. These results indicate that listeners can influence how many sound sources they perceive, as reflected in neural responses that track both the input and its perceptual organization.

**Significance Statement:** Can we consciously influence our perception of the external world? We address this question using sound sequences that can be heard either as coming from a single source, or as two distinct auditory streams. Listeners reported spontaneous changes in their perception between these two interpretations while we recorded neural activity to identify signatures of such integration and segregation. They also indicated that they could, to some extent, choose between these alternatives. This claim was supported by corresponding changes in responses in auditory cortex. By linking neural and behavioral correlates of perception we demonstrate that the number of objects we perceive can depend not only on the physical attributes of our environment, but also on how we intend to experience it.

## Introduction

For us to make sense of our environment, the brain must determine which elements of energy arriving at the sensory organs arise from the same source and should therefore be perceptually grouped. In audition, the less rapidly that sequential sounds change in one or more physical quantities, such as frequency, intensity, or spatial location, the more likely they are to be integrated and represented as a single perceptual object or stream (Moore & Gockel, 2012; van Noorden, 1975). The processes that underlie integration and segregation are affected not only by these stimulus features but also by internal states of the listener, such as the degree to which they are attending to the sounds (Carlyon et al., 2001; Sussman et al., 2002; Snyder et al., 2006; Billig and Carlyon, 2015), and by whether the stimuli correspond to a familiar speaker (Johnsrude et al., 2013) or word (Billig et al., 2013). The extent to which observers can voluntarily influence how they perceptually organize the outside world is unclear and bears on questions of whether and how higher-level cognition can influence perception (Fodor, 1983; Pylyshyn, 1999; Firestone and Scholl, 2015; Gross, 2017; Lupyan, 2017).

A common stimulus for investigating auditory perceptual organization is a repeating pattern of pure tones of high (H) and low (L) frequencies, such as that shown in Figure 1A. For lower frequency separations and presentation rates the sounds tend to be heard as integrated in a single stream that forms a distinctive galloping rhythm. At greater frequency separations and presentation rates the H and L tones typically form two segregated streams (van Noorden, 1975). For a range of stimulus parameters, perception can alternate between the two percepts every few seconds, usually after a longer initial integrated phase (Carlyon et al., 2001; Denham et al., 2013; Pressnitzer & Hupé, 2006; Figure 1B).

**Figure 1.**
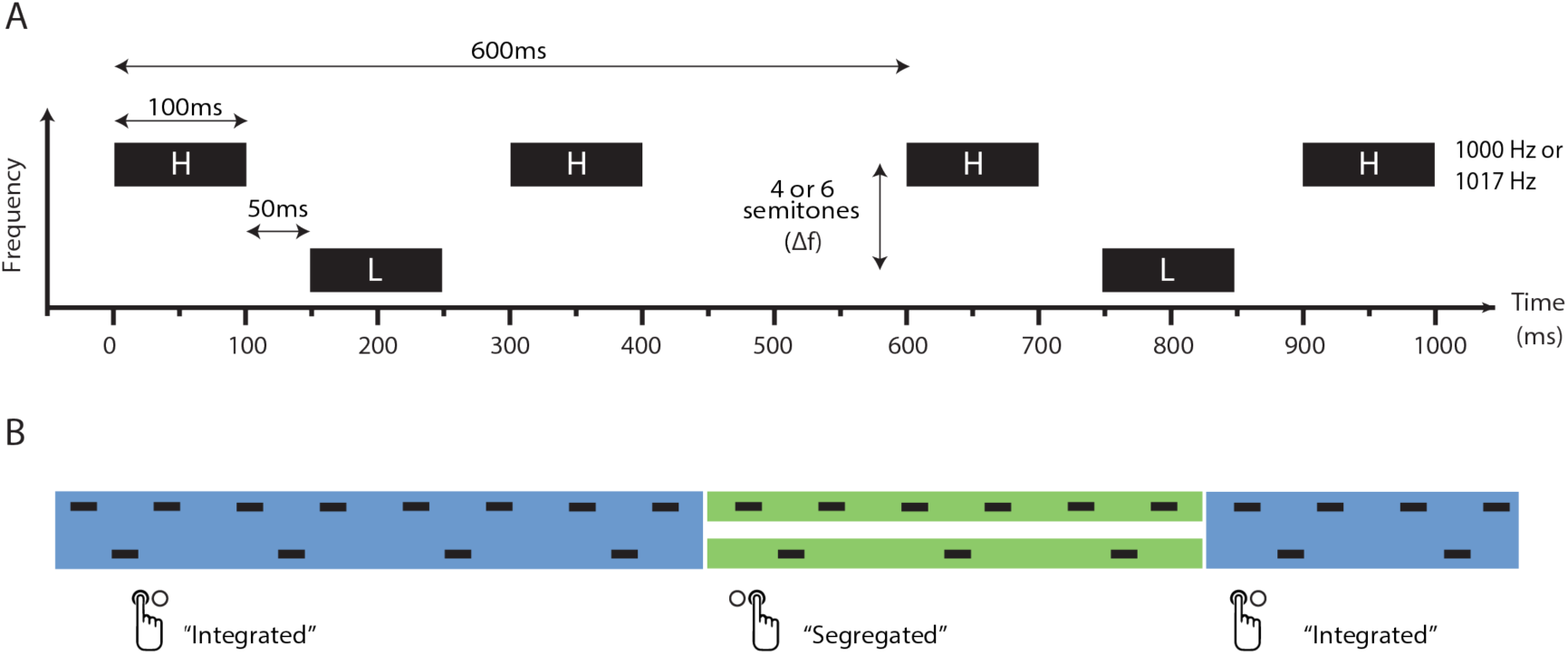
Stimulus parameters and percept reporting. (**A**) Two triplets of a stimulus sequence consisting of high (H) and low (L) tones with a frequency separation (Δf) of 4 or 6 semitones. The H tone frequency was 1000 Hz for soundbooth testing and 1017 Hz for testing with electro-/magneto-encephalography. (**B**) Illustrative changes in perceptual organization with corresponding button press reports. During integration (blue), H and L tones are perceived as belonging to a single pattern, whereas during segregation (green) they form two separate perceptual streams. Perception typically alternates every few seconds after a longer initial integrated phase.

For such ambiguous sequences, listeners report being able to exert a degree of control over hearing integration or segregation (van Noorden, 1975; Pressnitzer and Hupé, 2006; Micheyl and Oxenham, 2010; Farkas et al., 2016). However, subjective responses may be affected by post-perceptual processes and biases, such as shifts in decision criteria (Green and Swets, 1966) and attempts to meet the perceived aims of the experiment (Orne, 1962). To the extent that they vary with a listener’s percept, indirect behavioral or neural measures can bypass such issues. For example, several electro-and magneto-encephalography (EEG/MEG) studies have detected more positive auditory cortical responses approximately 60-100 ms following the onset of the middle tone in such triplets during reports of segregation compared to integration (Gutschalk et al., 2005; Hill et al., 2012; Szalárdy et al., 2013a). We argue that these objective neural measures can also shed light on the neural stages of processing that underlie any genuine effect of intention on perception.

Here we combine subjective and objective measures to demonstrate an effect of intention on perception, reflected in evoked responses in auditory cortex. To do so we measure neural activity with EEG/MEG as participants listen neutrally to HLH-sequences (Figure 1A) and report spontaneous changes in their perception (Figure 1B). We derive a univariate marker of perceptual organization in the auditory evoked field at the group level, but also make use of multiple temporal features in the neural response to train multivariate percept classifiers for each participant. We then study the relative occurrence of these neural signatures when participants actively try to promote integration (by attending to the whole pattern) or segregation (by attending exclusively to tones of one frequency). This allows us to establish whether their reports of successfully influencing their percept are supported by and reflected in stimulus-locked activity in auditory cortex, or are instead more likely to have a post-perceptual locus.

## Materials and Methods

### Participants

Data were collected in two separate experimental settings. Twenty-five participants took part in a sound booth (Setting 1), for the purposes of (a) ensuring that stimulus parameters gave rise to integrated and segregated percepts in approximately equal measure, and (b) screening participants before EEG/MEG recording to ensure that they could experience both percepts. Twenty-two of these participants also took part in the EEG/MEG lab (Setting 2), between 1 and 34 days later. Two further participants took part in Setting 2 only, after screening with an online test. All 27 participants across both settings were aged 18-40 (mean age = 28.56 years, 12 females), right-handed and reported no neurological or developmental disorders. They were recruited from the MRC Cognition and Brain Sciences Unit participant panel or by word of mouth, and were paid for their time. One participant, whose results were not qualitatively different from the remainder of the group, had a threshold of 30 dB HL at 1500 Hz in the left ear. All other participants had normal hearing (<25 dB HL pure tone thresholds over the range of the stimuli, 1000-2000 Hz). All experimental procedures were approved by the Cambridge Psychology Research Ethics Committee.

### Stimuli

Sequences of 250 HLH triplets were presented diotically, where H (high) and L (low) were 100-ms pure tones (Figure 1A). The frequency of the H tone was fixed at 1000 Hz (Setting 1) or 1017 Hz (Setting 2), except for the final H tone in the final triplet when it was 250 Hz. The choice of 1017 Hz in Setting 2 was to avoid possible contamination by harmonics of the 50-Hz line noise. The 250-Hz tone was low enough in frequency to be detectable on a low-pass filtered auxiliary channel of the MEG recording set-up in Setting 2, and was included to enable neural recordings to be time-locked to the stimulus. The L tone in a given sequence was lower than that of the H tone by an amount (Δf) of either four or six semitones (both settings). Silent intervals of 50 ms separated tones within a triplet, and silent intervals of 200 ms separated one triplet from the next, giving sequences of 150 s duration. These stimuli were chosen to match Experiment 2 of Gutschalk et al. (2005).

Filler stimuli lasting a total of 40 s were created to separate experimental sequences from each other. These consisted of 5,005 ms silence, followed by a 100-ms 250-Hz tone (a time-locking signal, with the same purpose as that described in the previous paragraph). This was followed by 1,900 ms of silence, then by 33 pure tones, each of 100 ms duration with 50 ms of silence between tones. The frequencies of these tones were selected at random from a log-rectangular distribution from 200-2000 Hz. Their purpose was to interfere with memory of the previous sequence in an effort to minimize context effects, such as those described by Snyder et al. (2009). The filler stimulus continued with 22,945 ms of silence, another 100-ms 250-Hz tone (to warn the participant that the next experimental sequence was about to begin) and a final 5,000 ms of silence. All tones in the experimental sequences and filler stimuli included 10-ms linear onset and offset ramps, and were generated digitally at a sample rate of 44100 Hz with 16-bit resolution.

### Experimental procedures

In Setting 1, participants were seated in a double-walled sound-insulated room and sounds were presented over Sennheiser HD650 headphones at a level of 55 dB SPL. In Setting 2, participants sat under the dewar of a VectorView system (Elekta Neuromag) while MEG and EEG activity was recorded (see “EEG and MEG acquisition and pre-processing” section for details of preparation and recording). In this setting, sounds were presented through tube headphones with silicone inserts at 50 dB above the participant’s 1000-Hz pure tone hearing threshold. Using their right hand, participants pressed one computer key (Setting 1) or button box button (Setting 2) when hearing an integrated, galloping triplet pattern, and another when hearing the tones segregate into two isochronous sequences (Figure 1B). The screen indicated their most recent response, which corresponded to their current percept. They were told to make a selection as soon as possible after the sequence began, and to make further responses whenever their percept changed. There were four conditions with different instructions. In “Neutral” sequences participants were instructed to let their perception take a natural course. In “Attempt Integration” sequences they tried to promote the integrated percept by attending to the whole pattern. In “Attempt Segregation” sequences they tried to promote the segregated percept by attending either to the H tones (“Attend High”) or the L tones (“Attend Low”).

The experiment consisted of two (Setting 1) or four (Setting 2) blocks. Each block contained five sequences: two Neutral, one Attempt Integration, one Attend High and one Attend Low. In Setting 1 the order of instruction conditions was the same in both blocks for a given participant; in Setting 2 this order was reversed for the final two blocks. The two Neutral sequences in a block never occurred consecutively, and Δf alternated between four and six semitones from sequence to sequence. An on-screen message specified the instruction prior to and throughout each trial. Response key/button mapping, and order of instruction and Δf conditions were balanced across participants. Participants relaxed between sequences and took breaks of at least a minute between blocks (while remaining under the dewar in Setting 2). During experimental trials in Setting 2 they were instructed to keep their eyes open and to maintain fixation on a cross in the centre of the screen, or elsewhere if more comfortable, to minimize alpha power and artefacts from eye movements. In Setting 2, participants’ head positions were checked at the start of each block, and their position adjusted (to minimize loss of MEG signal) if they had dropped by a centimetre or more. Testing lasted approximately 30 min in Setting 1 and 60 min in Setting 2.

Before the experiment, the concept of streaming was explained using HLH-patterns with Δf of 0, 5 and 12 semitones. Participants practiced reporting their percept while listening neutrally. They were then told that they may be able to influence their percept by attending either to the whole pattern or to one or other sets of tones; these conditions were also practiced. Participants were told that it was far more important to be honest and accurate in their responses than to be successful in their attempts to influence their percept. In Setting 2, practice and experimental blocks occurred after electrode preparation and head position digitization (described in the “Experimental design and statistical analysis - EEG and MEG acquisition and pre-processing” section). The two participants who had not taken part in Setting 1 completed an online training session to familiarize themselves with the stimuli and percept reporting process, and to practice trying to influence their percept. Instructions were repeated in person immediately prior to the experiment. Those participants who had taken part in Setting 1 more than a week previously also completed the online training as a refresher.

### EEG and MEG acquisition and pre-processing

Magnetic fields were recorded using a VectorView system (Elekta Neuromag) with one magnetometer and two orthogonal planar gradiometers at each of 102 locations. Electric potentials were recorded concurrently using seventy Ag-AgCl sensors arranged in the extended 10-10% configuration, fitted to the scalp using an electrode cap (Easycap) and referenced to an electrode on the nose, with a ground electrode on the right cheek. Head position was continuously monitored using five head position indicator (HPI) coils. Electro-cardiographic (ECG) and horizontal and vertical electro-oculographic (EOG) activity was recorded with three pairs of electrodes. The positions of the EEG sensors, HPI coils and approximately 100 additional head points were digitized with a 3D digitizer (Fastrak Polhemus), relative to three anatomical fiducial points (the nasion and both pre-auricular points). Data were acquired with a sampling rate of 1000 Hz and a high-pass filter of 0.01 Hz. For the magnetometer and gradiometer recordings, the temporal extension of Signal Space Separation in MaxFilter was used to identify bad channels, suppress noise sources, and compensate for head movement. For all sensor types, additional noisy channels were identified and excluded for each participant based on observations during recording and offline visual inspection, as were recording segments containing SQUID jumps, channel pops, and muscle activity. Line noise at 50 Hz and its harmonics was removed using adaptive multitaper regression implemented in the EEGLAB plugin CleanLine, after which all activity was downsampled to 250 Hz. Independent components analysis (ICA) was performed in EEGLAB using the Infomax routine (with sub-Gaussian components included) on a version of the data that had been high-pass filtered at 0.5 Hz (6 dB cut-off, 1 Hz transition band, FIR windowed sinc filter) to impose the stationarity assumed by ICA. EEG channels were considerably noisier than magnetometers and gradiometers, and did not improve the quality of the decomposition. They were therefore discarded, and subsequent analyses were restricted to magnetometers and gradiometers only. Components corresponding to eye blinks/movements and cardiac artefacts were identified and projected out of another copy of the data that had been low-pass filtered at 30 Hz (6 dB cut-off, 6.667 Hz transition band, FIR windowed sinc filter) and high-pass filtered at 0.278 Hz (6 dB cut-off, 0.556 Hz transition band, FIR windowed sinc filter). This high-pass filter was selected for reasons explained in the next paragraph.

The resulting data were divided into 600-ms epochs, each beginning at the start of an HLH-triplet. Epochs beginning less than 1500 ms after a button press (or the start of the sequence) or ending less than 1500 ms before a button press (or the end of the sequence) were excluded from analyses. This minimized neural and muscular activity related to movement, and removed periods around transitions when the reported percept was least likely to be reliable. Baseline correction was not performed due to the repeating nature of the stimulus precluding a sufficient silent period between triplets, which meant that neural responses from one epoch were likely to carry over to the next. Due to the exclusion of epochs close to reported perceptual switches, any such influence should arise solely from triplets with the same (reported) perceptual state, and epoch time can therefore be thought of as circular (see Hill et al. (2012) for a similar approach). The high-pass filter of 0.278 Hz corresponds to a 3600-ms time period, the shortest possible interval between retained epochs corresponding to different perceptual reports. The relatively conservative approach of epoch rejection, necessary for tapping periods that were as perceptually stable as possible, led to a median retention rate of 58% (2900 epochs) per participant, comparable to that in Hill et al. (2012).

### Dipole fitting

Pairs of equivalent current dipoles were fitted to the magnetometer and gradiometer data for each participant separately, using the VB-ECD approach in SPM12. Reconstructions made use of single shell forward models based on participant-specific T1-weighted structural MRI scans. Sensor positions were projected onto each participant’s MRI by minimizing the sum of squared differences between the digitized fiducials and MRI scan fiducials, and between the digitized head shape and the individual scalp mesh. The VB-ECD routine uses a variational Bayes approach to iteratively optimize location and orientation parameters of fitted dipoles. The midpoints of each hemisphere’s Heschl’s gyrus were used as soft location priors, with no priors for dipole orientation. Fitting was performed separately for magnetometer and gradiometer data, using the mean activity in the 24-ms window centered on the first prominent turning point in the sensor space waveform (peaking 40-110 ms after triplet onset), over all epochs. The dipole pair that accounted for the most variance in sensor data out of 20 iterations of the fitting process was selected for each participant and each sensor type. Using these dipoles as spatial filters, further analyses were conducted on the hemisphere-specific source waveforms, and on the mean waveform across hemispheres. As the polarity of reconstructed waveforms depends on the orientation of the sources with respect to individual anatomy, each participant’s source waveforms were inspected and inverted as necessary such that the first prominent turning point (peaking 40-110 ms after triplet onset) was a local maximum. All results were comparable across magnetometers and gradiometers, and are reported for gradiometers only. Fitted dipole pairs accounted for a mean of 91.9% (standard deviation 4.2%) of the variance in the sensor recordings over the fitting window and were located in or close to Heschl’s gyrus for all hemispheres (mean MNI coordinates [+/−49 −21 3], standard deviation 6 mm).

To verify that our findings were not dependent on the use of location priors in Heschl’s gyrus, we performed a separate set of analyses, selecting for each participant the neural component from the ICA that had the maximum back-projected power in the evoked response. Dipoles fitted to these components also had a mean location in Heschl’s gyrus, and the reconstructed source waveforms showed qualitatively similar results to those described below. Although for some participants these reconstructed sources were located in regions remote from auditory cortex, their locations were not consistent across participants and not considered further.

### Experimental design and statistical analysis

#### Sample size justification

No published research has used the same approach to test for intention effects with the same stimuli, however effect sizes for two relevant findings can be estimated from previous studies: (a) an intention effect on behavioral streaming measures of η^2^=.70 (Pressnitzer & Hupé, 2006) and (b) a percept effect on MEG evoked responses of η^2^=.43 (Gutschalk et al., 2005). Of the two effects, the latter would require the largest sample size to detect, namely 18 participants for 90% power. We tested 24 participants in the MEG setting; this allowed for drop-outs and accounted for possible over-estimation of effect sizes due to unreported null findings.

#### Behavioral analyses

As there were no significant differences in the mean percentage of segregation reported across the sound booth and EEG/MEG lab settings (*t*(15)=1.38, *p*=.189, *d*=0.23, 95% CI [-0.20 0.04], tested on the 16 participants with no conditions in either setting in which the percentage of segregation was 0 or 100), behavioral data were combined across the two settings. Of the 27 participants, two (both tested only in Setting 1) were excluded from behavioral data analyses. Both had at least one Δf × instruction condition with no sequences that met the following criteria: (i) the first reported phase was integrated (ii) at least two completed subsequent phases were reported. These criteria were necessary to allow separate analysis and comparison of the duration of initial-integrated, subsequent-integrated and segregated phases. Percentages of segregation for all remaining participants were logit-transformed and phase durations were log-transformed before being submitted to repeated-measures ANOVAs for analysis as a function of Δf and instruction. These transformations typically produced data with normally distributed residuals. When this was not the case, non-parametric tests were also conducted; these gave rise to the same qualitative pattern of results and are not reported separately. Mean percentages/durations were calculated on the transformed scale, then converted back to percentages/seconds for reporting. Null hypothesis significance testing was applied, with an alpha value of .05. Degrees of freedom were adjusted for asphericity as appropriate using the Huynh-Feldt correction (uncorrected degrees of freedom are reported for clarity).

#### Univariate neural analyses

Epochs in the Neutral condition were averaged for each combination of Δf and reported percept, for each participant. To maximize power, epochs occurring prior to the first percept report were labelled as integrated, and the first integrated phase of a sequence was not considered separately from the remaining integrated phases. The exclusion of any epochs before the first segregated report (in line with some researchers’ suggestions to treat these separately (Denham et al., 2013)) led to qualitatively similar results. One participant, who had only five valid epochs in one Δf × percept cell in the Neutral condition, was excluded from subsequent analyses of neural data. All other participants had at least 55 valid epochs per Δf × percept cell in the Neutral condition (the mean across participants of number of epochs in smallest cell was 145). To assess neural activity as a function of percept without stimulus confounds, the timecourses of the two Δf conditions were averaged within each percept before statistical analysis. Percept differences were similarly partialled out of analyses of neural activity as a function of Δf.

Statistical differences between percepts and frequency separations in the Neutral condition were assessed using a cluster-based permutation method (Maris and Oostenveld, 2007). Within-participants *t*-tests were conducted at each timepoint, and the largest contiguous cluster of values all exceeding a critical *t*-value (corresponding to an alpha value of .05) was selected for further analysis. Cluster significance was assessed by comparison to a null distribution generated by randomly permuting the labels of condition averages 1000 times within each participant, and using an alpha value of .05.

Epochs in the Attempt Integration, Attempt Segregation (Attend High), and Attempt Segregation (Attend Low) conditions were averaged within each Δf, without regard to reported percept. Differences over the temporal cluster of interest from the Neutral condition were derived for each of the following contrasts: (a) ½ * Attempt Segregation (Attend High) + ½ * Attempt Segregation (Attend Low) – Attempt Integration, (b) Attempt Segregation (Attend High) – Attempt Integration, (c) Attempt Segregation (Attend Low) – Attempt Integration, (d) Attempt Segregation (Attend High) – Attempt Segregation (Attend Low). In all cases, the two Δf conditions were given equal weight. Paired *t*-tests were conducted on these differences.

To test whether effects of intention on univariate neural responses in the non-Neutral conditions were as large as would be expected based on perceptual reports, and under the assumption that the neural signature of percept in the Neutral condition also applied in the non-Neutral conditions, the following calculations were made. The percentages of each percept reported in each non-Neutral condition for each Δf and participant were applied to the relevant mean neural response from the Neutral condition. Simulated and observed values were compared using a paired *t*-test, with an alpha value of .05.

#### Multivariate neural analyses

Epochs were labelled and participants excluded as outlined in the “Experimental design and statistical analysis – Univariate neural analyses” section. Support Vector Machines (SVMs) with linear kernels were trained to classify integrated versus segregated epochs in the Neutral condition for each Δf and participant, using an adapted version of the DDTBOX package (Bode et al., 2017) in MATLAB. To ensure that the classifiers were unbiased, random sub-sampling within each SVM was used to match the number of epochs across classes. Five-fold cross-validation was applied, and the subsampling and cross-validation process was repeated 100 times. Features were the standardized values of the neural response at the 150 sampled time points of each 600-ms epoch (arising from the 250 Hz sampling frequency), and the cost parameter (C) was set as 1. Classifier performance in the Neutral condition was assessed for each participant by comparing classified versus actual labels and averaging the percent correct over the 5x100 = 500 iterations, and over Δf conditions. Group classification accuracy was tested against the 50% chance level using a *t*-test with an alpha value of .05. Feature weights were obtained from the SVM training functions, and corrected using the method of Haufe et al. (2014), which removes strongly weighted but theoretically irrelevant noise features. These were normalized across participants then averaged over Δf for plotting. The 500 trained SVMs for each Δf and participant were also used to classify all epochs in the non-Neutral conditions, regardless of percept report. The percentage classified as segregated was compared across non-Neutral conditions using within-participants *t*-tests with an alpha value of .05, for the same contrasts as outlined in the “Univariate neural analyses” section.

To test whether task-related differences in the percentage of epochs classified as segregated was as high as would be expected based on subjective reports, it was necessary to take into account the accuracy of the trained classifiers in the Neutral condition. The percentage of reports of segregation for each participant, frequency separation, and task was multiplied by (Neutral classification accuracy – 50)/50 (i.e. the Neutral classification accuracy above chance, as a proportion from −1 to 1). The expected task-related difference in the percentage of epochs classified as segregated was derived for each participant and frequency separation, by taking the average of these adjusted percentages of segregated reports over the two Attempt Segregation conditions and subtracting the adjusted percentage of segregated reports in the Attempt Integration condition. These expected difference values were then averaged over frequency separations, and compared to the observed differences using a paired *t*-test, with an alpha value of .05.

## Results

### Behavioral results

As shown in Figure 2A, segregation was reported for a greater proportion of time for the larger than for the smaller Δf for all tasks. This arose from a combination of shorter initial integrated phases (Figure 2B), shorter subsequent integrated phases (Figure 2C), and longer segregated phases (Figure 2D). All of these effects were statistically significant (Figure 2A: *F*(1,24) = 34.89, *p*<.001, η^2^_p_=.59, 95% CI [.48 .72]. Figure 2B: *F*(1,24)=46.58, *p*<.001, η^2^_p_=.66, 95% CI [.58 .77]. Figure 2C: *F*(1,24)=15.28, *p*<.001, η^2^_p_=.39, 95% CI [.18 .58]. Figure 2D: *F*(1,24)=11.28, *p*<.001, η^2^_p_=.32, 95% CI [.09 .58]).

**Figure 2.**
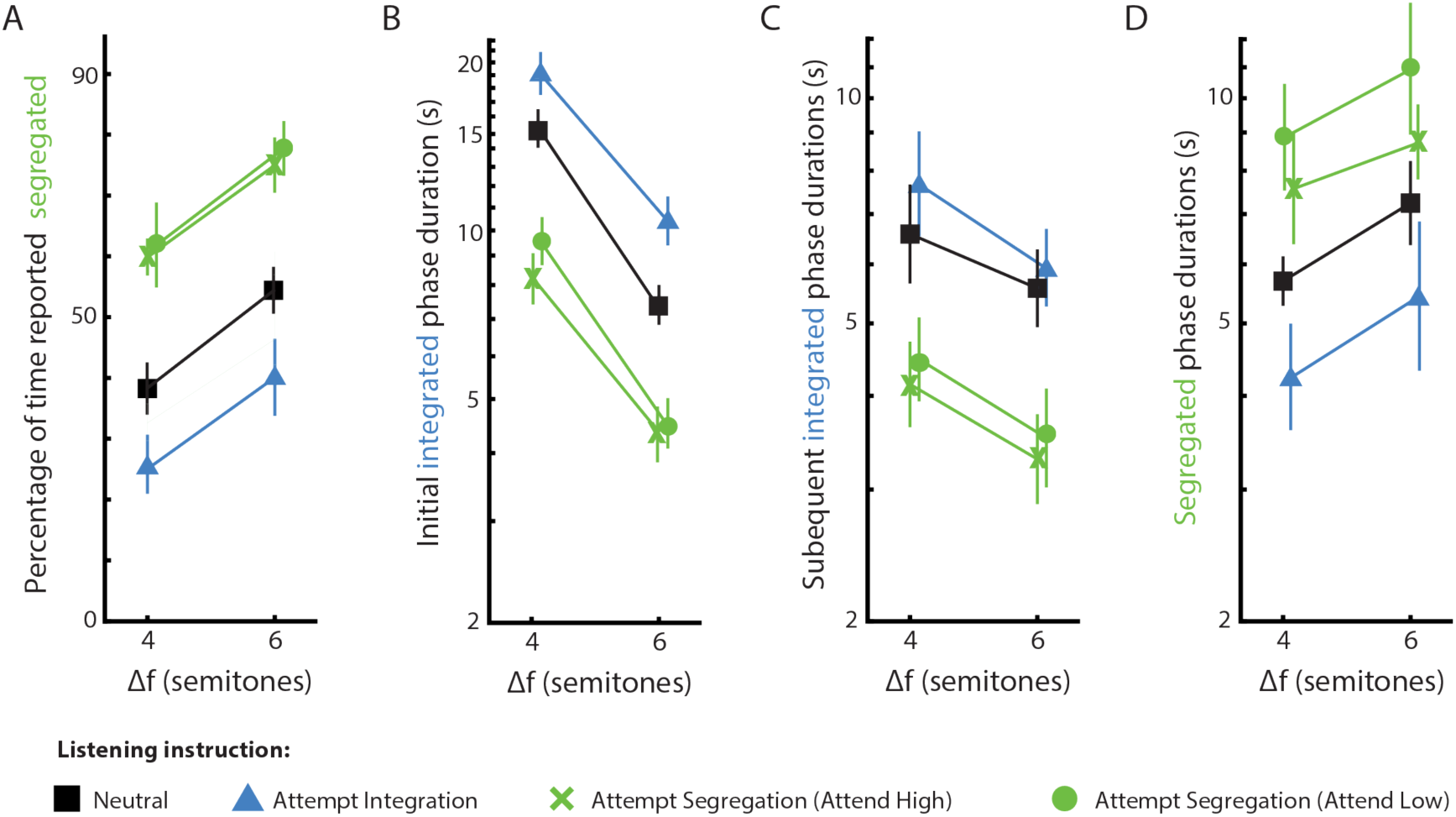
Behavioral analyses. Effects of frequency separation (Δf) and task on (**A**) the percentage of time reporting segregation, (**B**) the duration of initial integrated phases, (**C**) the duration of subsequent integrated phases, and (**D**) the duration of segregated phases. Phase durations are plotted on a log scale. Black squares: listen neutrally, blue triangles: attempt integration, green crosses: attempt segregation by attending to high tones, green circles: attempt segregation by attending to low tones. Error bars: 95% within-participants confidence intervals.

Importantly, the percentage of time each percept was reported was also affected by the task instructions (*F*(1,24)=51.55, *p*<.001, η^2^_p_=.68, 95% CI [.58 .80]; Figure 2A). This effect was reflected in extended phases of the intended percept (although not to a significant extent for non-initial integrated phases) and shortened phases of the unintended percept, in comparison to the Neutral condition (Figure 2B, 2C, 2D; see Table 1 for statistics). Focusing on tones of a single frequency to promote segregation had a larger effect on the percentage of time hearing segregation than trying to hold the three tones in a triplet together (the black lines are closer to the blue lines than to the green lines in Figure 2A; *t*(24)=3.03, *p*=.006, *d*=0.67, 95% CI [0.17 1.17]).

**Table 1.**
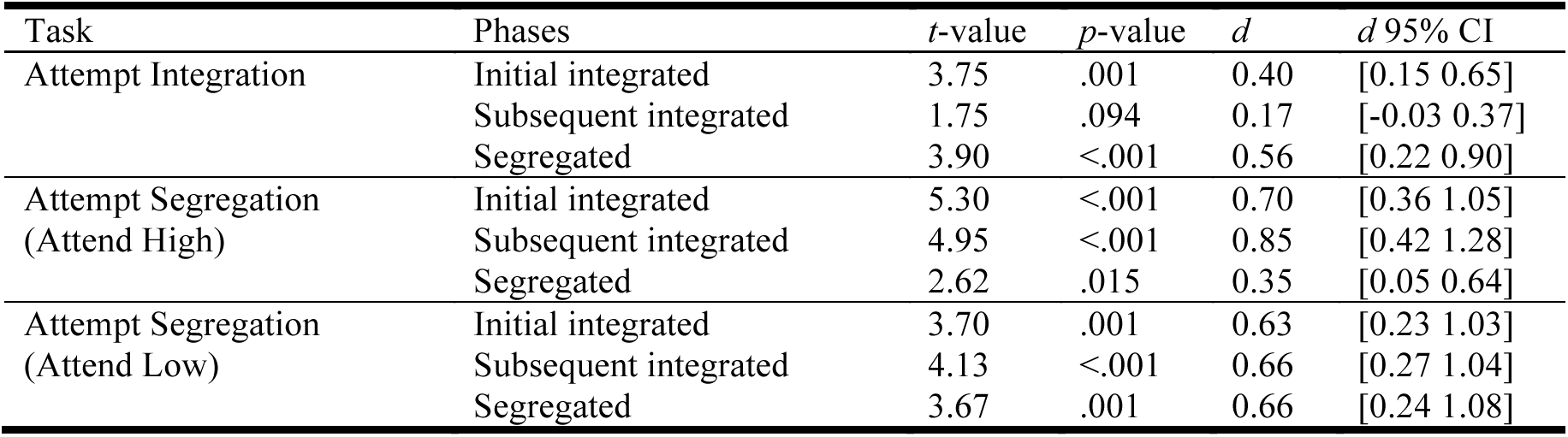
Statistics for effects of intention on phase durations (compared to the Neutral condition)

However, there was no effect of attending to the high versus the low tones during segregated listening (*t*(24)=1.29, *p*=.208, *d*=0.27, 95% CI [-0.15 0.69]).

### Univariate neural results

Neural responses time-locked to the onset of each HLH-triplet were extracted for each percept in the Neutral condition independent of Δf (Figure 3A), averaging over the dipoles in bilateral auditory cortices (Figure 3B). A univariate analysis revealed a time window 216-288 ms post triplet onset (66-138 ms post L tone onset) during which epochs reported as segregated evoked a significantly more positive response than those reported as integrated, independent of Δf (cluster-based permutation test, *p*=.001; window-specific test, *t*(22)=4.18, *p*<.001, *d*=0.32, 95% CI [0.12 0.52]; Figure 3C). When based on single dipoles, the size of the percept effect in this time window did not differ between the left and right hemispheres (*t*(22)=1.55, *p*=.135, *d*=0.35, 95% CI [-0.13 0.83]).

**Figure 3.**
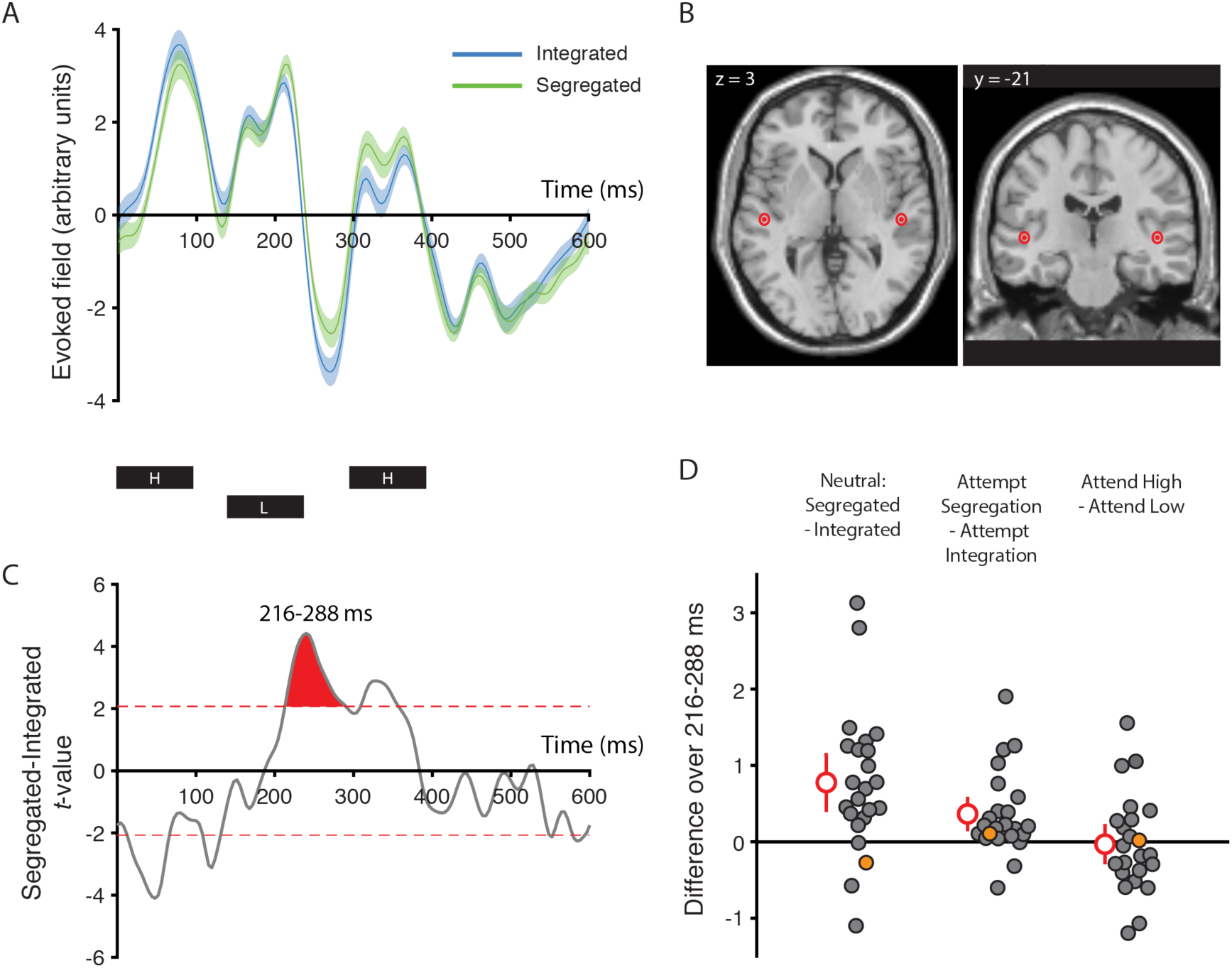
Univariate neural analyses. (**A**) Group timecourse (mean and 95% within-participants confidence intervals) of neural activity for integrated (blue) and segregated (green) reports in the Neutral condition, across frequency separations. Activity is projected through a spatial filter based on dipoles in bilateral auditory cortex fitted to the sensor data separately for each participant. The timing of each tone in the triplet is indicated below the plot. (**B**) Mean and 95% within-participants confidence interval of the fitted dipole locations for the activity in Figure 3A. Sources are shown on a template brain, with coordinates in MNI space. Mean reconstructed sources lie in bilateral posteromedial Heschl’s gyrus. (**C**) *t*-values for the Neutral Segregated minus Integrated group difference wave, across frequency separations. Dashed red lines indicate the critical *t*-values at *p*=.05, and the shaded red area represents the largest supra-threshold cluster. (**D**) Differences in neural activity averaged over the time window of interest for Neutral Segregated minus Integrated (left), Attempt Segregation minus Attempt Integration (middle), and Attend High minus Attend Low (right), across frequency separations. Filled circles correspond to individual participants, with mean and 95% confidence intervals shown in red. The orange circle represents a single participant also highlighted in Figures 4B, 4C, and 4D, for comparison of results across univariate and multivariate approaches.

**Figure 4.**
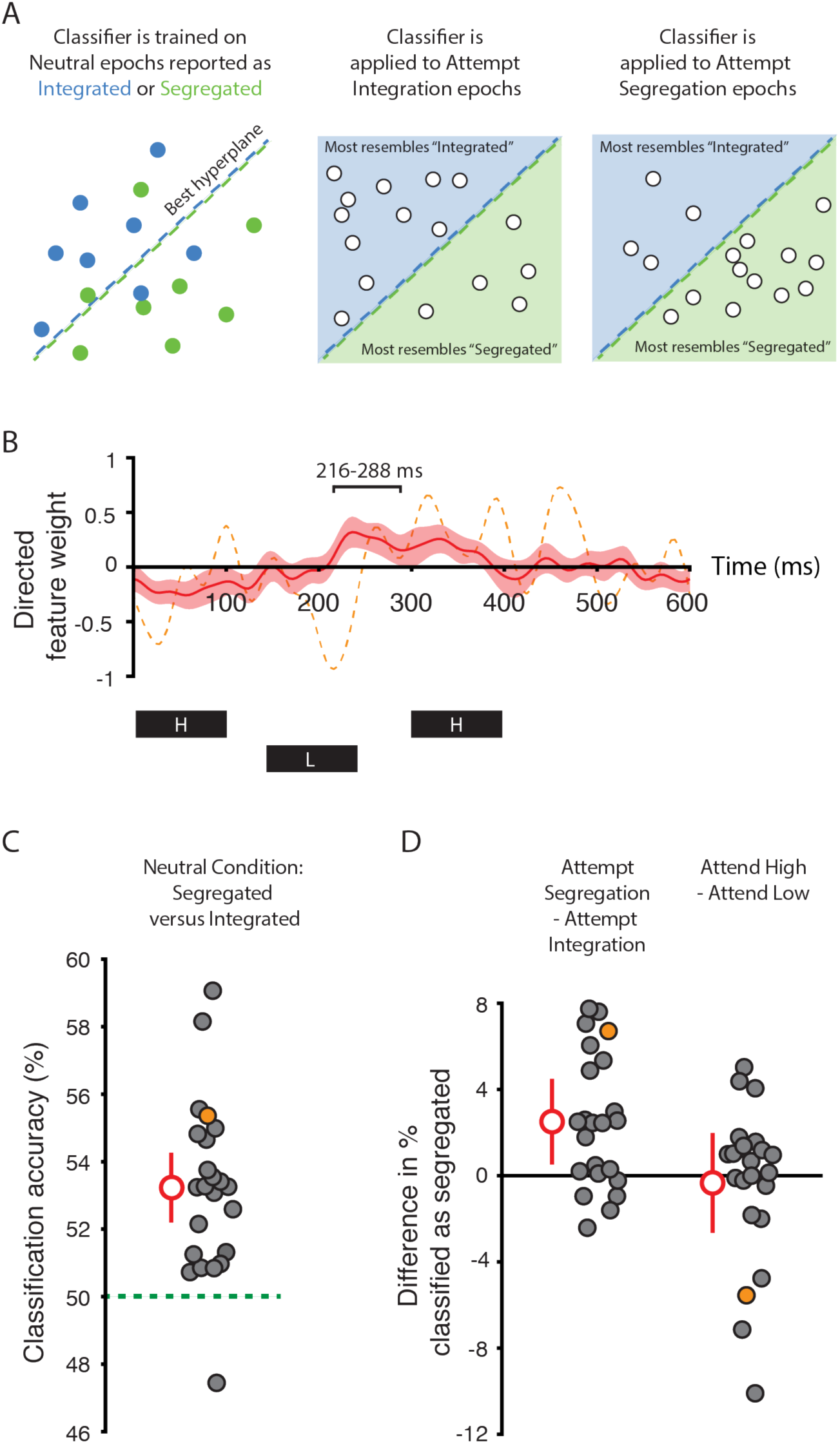
Multivariate neural analyses. (**A**) Schematic illustration of the classification approach. Linear Support Vector Machines (SVMs) are trained for each participant and frequency separation in the Neutral condition (left panel) to find the hyperplane (dashed line) that optimally separates epochs reported as integrated (blue circles) and segregated (green circles). The SVMs are then applied to epochs (white circles) in the Attempt Integration (middle panel) and Attempt Segregation (right panel) conditions. (**B**) Directed feature weights for classification in the Neutral condition, across frequency separations. A positive weight at a given timepoint reflects that a more positive neural response contributes to a segregated classification. The mean across the group is plotted in red with 95% confidence intervals in pink. One peak in the mean trace lies in the significant 216-288 ms window from the univariate analysis. However, classification can make use of different features (timepoints) for different participants. The dashed orange trace corresponds to one participant whose neural activity is dissimilar to the group mean (both for feature weights plotted here, and raw activity in the 216-288 ms window shown in Figure 3D, left) and whose data did not contribute to the univariate effect of intention (Figure 3D, middle). Perception can nonetheless be decoded for this participant (Figure 4C), contributing to the multivariate effect of intention (Figure 4D, left). The timing of each tone in the triplet is indicated below the plot. (**C**) Classification accuracy for Segregated versus Integrated epochs in the Neutral condition, across frequency separations. Filled circles correspond to individual participants, with mean and 95% confidence intervals shown in red. The orange circle represents a single participant also highlighted in Figures 4B, 4C, and 4D, for comparison of results across univariate and multivariate approaches, as described above. Chance classification accuracy is at 50% (green dashed line). (**D**) Differences in percentage of epochs classified as segregated for Attempt Segregation minus Attempt Integration (left), and Attend High minus Attend Low (right), across frequency separations. Filled circles correspond to individual participants, with mean and 95% confidence intervals shown in red. The orange circle represents a single participant also highlighted in Figures 4B, 4C, and 4D, for comparison of results across univariate and multivariate approaches.

The effect of intention on the neural response in this window was determined by subtracting the mean over all epochs during attempts at integration from the mean over all epochs during attempts at segregation, regardless of reported percept. The group difference was significantly greater than zero (*t*(22)=3.14, *p*=.005, *d*=0.17, 95% CI [0.05 0.29]; Figure 3D, middle), paralleling the percept comparison in the Neutral condition (Figure 3D, left) and supporting participants’ reports that they heard more segregation when they tried to do so than when they tried to hear integration. This effect is unlikely to be driven by attention-related modulations of neural responses to particular tones independent of perceptual organization; the conditions in which attention was focused on the H or the L tones did not differ significantly from each other (*t*(22)=0.23, *p*=.819, *d*=0.01, 95% CI [-0.11 0.14]; Figure 3D, right). Importantly, there was also no evidence for a residual response bias; the magnitude of the neural difference in the non-Neutral conditions was similar to that expected if all reports in those conditions were accurate (*t*(22)=0.38, *p*=.710, *d*=0.08, 95% CI [−0.35 0.51]).

### Multivariate neural results

The difference waveform in the Neutral condition (Figure 3C) indicated that multiple time windows might be informative in distinguishing between integrated and segregated percepts, beyond the 216-288 ms window determined from the univariate analysis. To make use of information across the entire epoch, we sought multivariate temporal patterns that distinguished between integrated and segregated percepts at a single-trial level, and which were allowed to vary across participants. Linear Support Vector Machines (SVMs) trained for each Δf and participant (Figure 4A) achieved classification accuracy significantly above chance (*t*(22)=6.11, *p*<.001, *d*=1.77, 95% CI [1.17 2.86]; Figure 4C); this was driven by responses in multiple time windows, including that identified in the univariate analysis (Figure 4B). When based on single dipoles, classifier performance did not differ between the left and right hemispheres (*t*(22)=1.02, *p*=.321, *d*=0.22, 95% CI [-0.21 0.68]).

The SVMs trained on Neutral epochs were then used to classify epochs in the other conditions. Paralleling the univariate results, a greater percentage of epochs were classified as segregated when participants attempted segregation than when they tried to integrate the sounds (*t*(22)=3.87, *p*<.001, *d*=1.12, 95% CI [0.63 1.77]; Figure 4D, left). Again, this was not driven by epochs in which tones of one particular frequency were attended; the percentage of epochs classified as segregated was similar whether participants attended to high or low tones (*t*(22)=0.45, *p*=.657, *d*=0.13, 95% CI [-0.59 0.63]; Figure 4D, right).

The task-related difference in the percentage of epochs classified as segregated (mean 2.5%) was more than an order of magnitude smaller than the difference in reported proportions (mean 36.6%). This discrepancy was due to non-perfect classifier performance; although accuracy was above chance (50%), the mean was only 53.2% and the maximum across participants 59.1%. After taking into account the accuracy of each classifier, the task-related difference in the percentage of epochs classified as segregated was no different from that expected if all percept reports in the non-Neutral conditions were accurate (*t*(22)=0.55, *p*=.586, *d*=0.08, 95% CI [-0.24 0.42]). In line with the univariate analysis, there was therefore no evidence for a residual response bias.

The effect of intention determined by the multivariate analysis was larger and more reliable than that from the univariate analysis. The more flexible approach was able to exploit the data of participants whose neural activity did not align with the group percept signature in the 216-288 ms time window. For example, one participant’s percept in the Neutral condition could be decoded above chance based on the activity at a range of timepoints, including an effect in the opposite direction from that of the group around 216 ms post triplet onset (Figure 4B, dashed orange trace; Figure 4C, orange circle).

## Discussion

Our findings demonstrate that listeners can exert intentional control over how many objects they perceive in an ambiguous auditory scene. Differences in auditory cortical responses during attempts to hear repeating patterns of pure tones as an integrated whole versus segregated streams were consistent with signatures of these percepts obtained during a neutral listening condition. These differences supported listeners’ subjective reports that they could, to some extent, “hear what they want to hear”.

### Indexing low-level perception

We argue that the activity measured during neutral listening relates to the percept rather than to decisions made during the process of reporting it. The inherent uncertainty of localization based on MEG precludes ascribing a primary versus non-primary auditory cortical locus; our source reconstruction appears consistent with either of these. However, it seems unlikely that post-perceptual decision-related activity would originate from auditory regions and be so consistently timed from the onset of each stimulus. Furthermore, we excluded epochs surrounding button presses to minimize the contribution of activity relating to motor planning or execution. We therefore take the neutral neural signature to reflect perceptual experience. The use of bistable stimuli to probe perception also avoided acoustic confounds. Although we presented stimuli with two different frequency separations, leading to different reported proportions of segregation (c.f. Gutschalk et al., 2005), the key comparisons of neural activity were between alternative percepts of identical sounds.

Our interpretation of activity in the non-Neutral conditions assumes that the neural response carried more information about perception than about the instructions themselves, which differed in terms of how listeners were to attend to the sounds. Selective attention is known to affect the evoked response to tones even when perceptual organization is stable (Hillyard et al., 1973; Näätänen et al., 1978). Such modulations would presumably be maximally different across the two sub-conditions in which participants attended exclusively to either the H or L tones, rather than between one of these sub-conditions and the case when listeners attended to all of the tones. However, we found no difference for attention to the H versus the L tones over the time window of interest in the univariate analysis, nor in the percentage of epochs classified as segregated in the multivariate analysis. We therefore argue that attention alone (without concomitant changes in perceptual organization) cannot account for the observed neural effects.

Another important feature of our design was the simultaneous collection of percept reports and neural data, allowing us to draw direct associations between the two. Some previous studies have inferred integration or segregation using measures sensitive to stimulus manipulations that also affect perceptual organization, such as the mismatch negativity (Sussman et al., 1999; Winkler et al., 2006; Carlyon et al., 2010) or performance on a deviant detection task (Carlyon et al., 2010; Micheyl and Oxenham, 2010; Billig et al., 2013; Spielmann et al., 2014). However such measures are influenced by additional factors (Divenyi and Danner, 1977; Spielmann et al., 2013, 2014; Sussman et al., 2013; Szalárdy et al., 2013b), and the degree to which they, in isolation, can provide a reliable indication of perceptual organization over the course of sustained bistable stimulation is unclear.

### Implications for auditory scene analysis

The more positive response for segregation compared to integration from 66-138 ms after the onset of the L tone was consistent with previous findings (Gutschalk et al., 2005; Hill et al., 2012; Szalárdy et al., 2013a). It may in part reflect an increased P1m response to the L tone during segregation, due to a release from adaptation by responses to the previous H tone as neuronal receptive fields narrow and segregation occurs (Fishman et al., 2001; Gutschalk and Dykstra, 2014). However, our results do not depend on this interpretation; given the continuous stimulation paradigm it is not clear how the observed differences relate to responses to individual tones. Furthermore, our participant-specific classification analysis indicated that this time window was not the most diagnostic of percept for all individuals. Variability across listeners may arise from distinct listening strategies, or reflect differences in how multiple components from repeated sounds summate to an aggregate measured signal. Multivariate techniques such as representational similarity analysis have provided insight into the fine spatial patterns representing stimulus information in the brain (Haxby et al., 2001; Kriegeskorte et al., 2008). Here we applied a different form of multivariate analysis - classification in the temporal domain - to reveal individualized percept-specific patterns in neural activity (see also Wilbertz et al., 2017, Reichert et al., 2014, for classification of bistable visual perception).

We observed effects of percept and intention when analyzing responses generated by neural sources in bilateral auditory cortex. Functional magnetic resonance imaging has also revealed greater responses in precuneus and right intraparietal sulcus during segregation compared to integration (Cusack, 2005; Hill et al., 2011). Our analysis of precisely stimulus-locked responses would have been insensitive to more temporally diffuse effects that such studies may have tapped. Further evidence for the involvement in streaming of a network beyond auditory cortex comes from activity during perceptual reversals (as opposed to during stable periods of integration or segregation) in inferior colliculus, thalamus, insula, supramarginal gyrus, and cerebellum (Kashino and Kondo, 2012; Kondo and Kashino, 2009; Schadwinkel and Gutschalk, 2010). How these regions support or reflect either spontaneous reversals or voluntary switches remains to be established.

A distinction has been drawn between primitive and schema-based processes of perceptual organization (Bregman, 1990). Primitive processes automatically partition a scene based on its physical properties, whereas schema-based processes select elements based on attention or prior knowledge. One might expect different neural instantiations of the outcomes of these processes; however, we found the same segregation signature regardless of whether listeners allowed their perception to take a natural course, deliberately attended to the H tones, or deliberately attended to the L tones. We argue that the neural realization of an auditory scene may not only consist of distinct representations of attended and unattended streams of differing fidelity (Mesgarani and Chang, 2012; Puvvada and Simon, 2017) but also mark whether any segregation has occurred at all (c.f. Gandras et al. 2017; Szalárdy, Winkler, et al. 2013).

We asked participants to try to influence their percept by attending either to a subset of the tones, or to all of them. The former approach may succeed by narrowing receptive fields of auditory cortical neurons such that different populations respond to the tones of each frequency (Fritz et al., 2007; Ahveninen et al., 2011), or by introducing a perceived loudness difference between H and L tones (van Noorden, 1975; Dai et al., 1991). In contrast, repeatedly shifting attention across frequencies may promote integration by disrupting these effects. The size of the change in reports from the Neutral to the Attempt Segregation condition was greater than that from the Neutral to the Attempt Integration condition. This was not the case in previous studies (Pressnitzer and Hupé, 2006; Micheyl and Oxenham, 2010), a fact that may reflect differences in stimuli, or in how instructions were interpreted. We also note that in our experiment, volitional control similarly affected reported durations of intended and unintended phases, whereas Pressnitzer and Hupé (2006) found that phases of unwanted percepts were curtailed to a greater degree than target phases were extended. Listeners in that study may have used additional strategies to shorten segregated phases, such as briefly diverting attention away from the tone sequence (Carlyon et al., 2003). Phase duration distributions have informed modelling of auditory scene analysis (Mill et al., 2013; Rankin et al., 2015) and prompted parallels to be drawn between different forms of bistability across sensory modalities (Pressnitzer and Hupé, 2006). We emphasize that the interaction between stimulus characteristics and high-level factors such as attention, which may differ across bistable phenomena, must be considered in general accounts of how the brain handles perceptual ambiguity (van Ee et al., 2005; Kogo et al., 2015).

### Summary

Auditory bistability offers a powerful means of understanding how cognitive states, such as listening goals, attention, and prior knowledge, influence perception, while controlling for stimulus differences. Linking subjective reports with neural measures on a trial-by-trial basis allows us to tap into low-level processes, as opposed to post-perceptual decisions. This method identifies signatures of perceptual experience in auditory cortex to demonstrate that listeners can not only use attention to enhance the representation of a subset of sounds, but also intentionally alter the number of distinct objects heard to make up the auditory scene.

## Acknowledgements

This research was supported by a Medical Research Council Doctoral Training Award Studentship for Alexander J. Billig and by Medical Research Council grant number MC-A060-5PQ70 for Robert P. Carlyon. Alexander J. Billig is currently affiliated with UCL Ear Institute, University College London, United Kingdom, and thanks Timothy D. Griffiths and Ingrid S. Johnsrude for continuing financial support while he wrote this paper.

